# Genetic Diversity Patterns and Domestication Origin of Soybean

**DOI:** 10.1101/369421

**Authors:** Soon-Chun Jeong, Jung-Kyung Moon, Soo-Kwon Park, Myung-Shin Kim, Kwanghee Lee, Soo Rang Lee, Namhee Jeong, Man Soo Choi, Namshin Kim, Sung-Taeg Kang, Euiho Park

## Abstract

Understanding diversity and evolution of a crop is an essential step to implement a strategy to expand its germplasm base for crop improvement research. Samples intensively collected from Korea, which is a small but central region in the distribution geography of soybean, were genotyped to provide sufficient data to underpin genome-wide population genetic questions. After removing natural hybrids and duplicated or redundant accessions, we obtained a non-redundant set comprising 1,957 domesticated and 1,079 wild accessions to perform population structure analyses. Our analysis demonstrates that while wild soybean germplasm will require additional sampling from diverse indigenous areas to expand the germplasm base, the current domesticated soybean germplasm is saturated in terms of genetic diversity. We then showed that our genome-wide polymorphism map enabled us to detect genetic loci underling flower color, seed-coat color, and domestication syndrome. A representative soybean set consisting of 194 accessions were divided into one domesticated subpopulation and four wild subpopulations that could be traced back to their geographic collection areas. Population genomics analyses suggested that the monophyletic group of domesticated soybeans was originated in eastern Japan. The results were further substantiated by a phylogenetic tree constructed from domestication-associated single nucleotide polymorphisms identified in this study.

## 1. Introduction

To fully capitalize on the vast reservoir of favorable alleles that control agronomic traits within wild and domesticated germplasm, extensive phenotyping and genotyping of germplasm collections are necessary. Soybean [*Glycine max* (L.) Merr.] is a major crop for dietary protein and oil worldwide. Several hundred soybean genomes have been resequenced (Lam et al. 2010; Chung et al. 2014; Zhou et al. 2015; Valliyodan et al. 2016) and three genome-wide high-density SNP arrays have been developed and used to genotype thousands of soybean accessions (Song et al. 2013; Lee et al. 2015; Wang et al. 2016). These data have been primarily used to compare the patterns of genetic variation between *G. max* and its wild progenitor (*G. soja* Siebold & Zucc.) to understand the history of soybean domestication and identify selective sweeps related to the domestication and improvement of soybeans. The data have also been used to identify loci controlling important agronomic traits, such as protein-and-oil and seed-weight traits (Hwang et al. 2014; Bandillo et al. 2015; Zhou et al. 2015). However, those studies have been limited to detecting or confirming the major genetic loci reported in previous genetic mapping studies using biparental populations. Further efforts will be required to implement genome-wide association studies (GWAS) (McCarthy et al. 2008) with higher statistical power and mapping resolution in soybean.

Soybean was domesticated ~5000 years ago from *G. soja*, its sympatric wild annual progenitor that is distributed throughout East Asia, including most of China, Korea, Japan, and part of Russia (Hymowitz 2004; Larson et al. 2014). Different regions of China have been proposed as a single center of soybean domestication on the basis of morphological, cytogenetic, and seed protein variation (Broich and Palmer 1981; Hymowitz and Kaizuma 1981; Hymowitz 2004). Multiple centers of domestication including the southern areas of Japan and China have also been proposed based on chloroplast sequence variation (Xu et al. 2002) and archaeological records (Lee et al. 2011). However, recent phylogenetic studies using whole-genome resequencing data clearly indicated a monophyletic nature of domesticated soybean (Lam et al. 2010; Chung et al. 2014; Zhou et al. 2015). Of late, molecular studies that used hundreds of markers and accessions have proposed different areas, such as the Yellow River of China (Li et al. 2010) and southern China (Guo et al. 2010), as a center of soybean domestication. Yet, two other studies suggested the domestication center as northern and central China using high-density SNP array data (Wang et al. 2016), or central China surrounding the Yellow River using specific-locus amplified fragment sequencing data (Han et al. 2016).

In most of the previous studies, accessions collected from the Korean Peninsula were underrepresented, although this region is a central region of wild soybean distribution. For example, in the recent genome-wide analyses reported by Wang et al. (2016) and Han et al. (2016), accessions collected from China accounted for 91.8% and 100% of the total samples, respectively. Here, we present an analysis of SNP genotype data from 2,824 domesticated, 1,360 wild, and 50 putative hybrid accessions as part of an effort to characterize the entire Korean indigenous soybean collection deposited in the country’s National Agrobiodiversity Center. We genotyped soybean accessions using the 180K Axiom® SoyaSNP array that was developed from soybean genome resequencing data (Lee et al. 2015). Our high-density SNP array data allowed us to evaluate levels of genetic diversity and patterns of population structure. We further attempted to detect genetic loci underlying soybean domestication and important agronomic traits, as well as provide a refined model of the evolutionary history of domesticated soybean.

## 2. Materials and Methods

### 2.1. Plant materials and SNP genotyping

The majority of the accessions were from the National Agrobiodiversity Center in Jeonju, Korea, with a small number of accessions provided by individual laboratories (Table S1). The National Agrobiodiversity Center collection consists of approximately 12,000 accessions of improved and landrace cultivars (*G. max*) and wild soybean (*G. soja*). The Korean germplasm collection substantially overlaps with those of other countries, particularly the United States. Most accessions collected from locations in other countries than Korea have been donated from the US National Genetic Resources Program. Notable exceptions were the 46 wild accessions from Japan, whose accession codes start with ‘B’. These were donated from National BioResource Project in Japan. As our primary goal was to characterize the indigenous soybean collection of Korea, we attempted as much as possible to genotype accessions unique to the Korean collection. At the same time, we analyzed representative sets of landrace accessions from China, North Korea, and Japan and approximately 400 improved lines, most of which are immediate descendants of ancestral lines of United States soybean cultivars (Gizlice et al. 1994), so that cultivated soybean from Korea could be assessed in the context of worldwide soybean germplasm pool (Table S2). Representative *G. soja* accessions from China, Russia, and Japan were also selected, allowing the the geographic distribution of wild soybean in each of these countries to be sampled. Initially, we planted approximately 5,000 domesticated and 2,400 wild soybean accessions, each of which contains approximately 90% of the accessions collected in Korea. After pure line selection by single seed descent was performed at least two times, DNA samples from approximately 4,400 diverse soybean germplasm lines were genotyped. However, because our SNP array data set ended up with smaller number of *G. soja* accessions from China than those from Korea and Japan, soybean population structure from the representative set was additionally assessed using genome resequencing data (downloaded from Figshare database, http://figshare.com/articles/Soybean_resequencing_project/1176133) from 45 *G. soja* accessions reported by Zhou et al. (2015).

DNA samples from the ~ 4,400 diverse soybean accessions were extracted from a single plant of each accession and were genotyped with the Axiom® SoyaSNP array containing 180,961 SNP sites (Lee et al. 2015). Of the lines genotyped, 4,234 with >97% sample call rate were selected for further analysis. SNPs were scored following the Axiom ® Genotyping Solution Data Analysis User Guide (http://www.affymetrix.com/) as described by Lee et al. (2015). Of the 180,961 SNPs, 170,223 were selected on the basis of the development and validation study. Missing data points in the 170,223 SNPs were imputed using BEAGLE 4.0 with default settings (Browning and Browning 2007). The 170,223 SNPs were then used to screen out duplicated and redundant accessions, leaving 3,036 non-redundant accessions. After the initial filtration, SNPs with heterozygous rate > 0.02 and minor allele frequency < 0.02 were discarded from the genotype data of the non-redundant accessions, leaving a total of 117,095 high quality SNPs for the further population analyses.

Phenotypic data used for GWAS were obtained primarily from field evaluations in the field at National Institute of Crop Science, Jeonju, Korea, in 2012 and 2013 (Table S1). The observed phenotype data were converted into binary data. The flower color phenotypes were divided into absence of color (white) or presence of colors ranging from light to dark purple. The seed coat color phenotypes were divided into absence of colors (yellow or green) or presence of colors ranging from brown to black. Domestication phenotypes were divided into presence (*G. max*) or absence (*G. soja*) of domestication.

### 2.2. Population structure and genetic diversity pattern analyses

Principal component analysis (PCA) was conducted to summarize the genetic structure and variation present in the soybean collection using smartpca function in Eigensoft v7.2 (Patterson et al. 2006; Price et al. 2006). We plotted the first three PCs. NJ trees were constructed by MEGA7 (Kumar et al. 2016) under the *p*-distances model. We used a model-based clustering method implemented in ADMIXTURE v1.23 (Alexander et al. 2009) to investigate the population structure of the soybean accessions. We determined the optimal *K*, the number of clusters based on the smallest cross-validation error calculated from v-fold cross-validation procedure. We plotted the membership coefficient using DISTRUCT (Rosenberg 2004). To investigate the level of genetic diversity maintained in soybean accessions, we calculated the nucleotide diversity (π) using VCFtools v 0.1.13 (Danecek et al. 2011). Genetic differentiation (Weir and Cockerham’s *F_ST_* (Weir and Cockerham 1984)) between *G. max* and each of the *G. soja* subpopulations was calculated using the VCFtools V0.1.13. Hierarchical analyses of molecular variance (AMOVA) in the whole soybean set and the representative soybean set were performed using ARLEQUIN v.3.5.2.2 (Excoffier and Lischer 2010). The significance of the values for *F_CT_* (difference among groups), *F_SC_* (difference among populations within groups), and *F_ST_* (difference among populations) was tested by 1023 permutations.

### 2.3. Genome-wide association studies

We conducted GWAS using PLINK 1.9 (Purcell et al. 2007). Flower color, seed-coat color, and domestication phenotype data were converted to binary data to perform a conditional logistic regression model analysis. Conditional specific SNPs were selected on the basis of minor allele frequencies of SNPs among the groups defined based on PCA analysis. A Bonferroni correction was used to control for the multiple testing problem by adjusting the alpha value from α = 0.01 to α= (0.01/117,095 SNPs) where 117,095 is the number of statistical tests conducted. Therefore, statistical significance of a SNP-trait association was set at 8.54e^−8^ (−log_10_ *P* = 7.06). Manhattan plots were produced using the qqman package (Turner 2014). To define linkage disequilibrium (LD) patterns, correlation coefficient of alleles (*r^2^*) were calculated for SNPs under the peak regions that exhibited significant association using Haploview 4.2 (Barrett et al. 2005). The confidence interval (CI) method of Gabriel et al. (2002) was used to identify LD blocks.

## 3. Results

### 3.1. Overall structure of the genotyped soybean germplasm population

Of the approximately 4,400 accessions genotyped, 4,234 exhibited >97% sample call rate. These were used as the total population for characterizing the Korean soybean germplasm (Table S1). The majority of this 4,234 set contained accessions from Korea (78.7% *G. max* and 91.5% *G. soja*) (Fig. 1A). The rest were *G. max* landrace and *G. soja* accessions from China, Russia, North Korea, and Japan and improved lines mostly developed in the United States (Table S2). To eliminate potential confounding effects exerted by hybrids in the comparison of wild and domesticated soybean populations (Vaughan et al. 2008; Wang et al. 2017), we first removed 50 putative hybrid accessions from among the 4,234 accessions (Fig. 1B; Table S2). In the field evaluation, most of these 50 accessions showed intermediate morphologies between domesticated soybean (*G. max*) and its wild relative (*G. soja*). Furthermore, principal component analysis (PCA) using 170,223 high-quality SNPs showed that the accessions were positioned between two large groups of *G. max* and *G. soja* (Fig. 1A). In further support of their suspected hybrid status, 50 accessions showed mixed wild or domesticated genome fractions ranging from 30 to 70% in *K* = 2 or 3 populations in the ADMIXTURE analysis (Fig. 1B).

**Figure 1.**
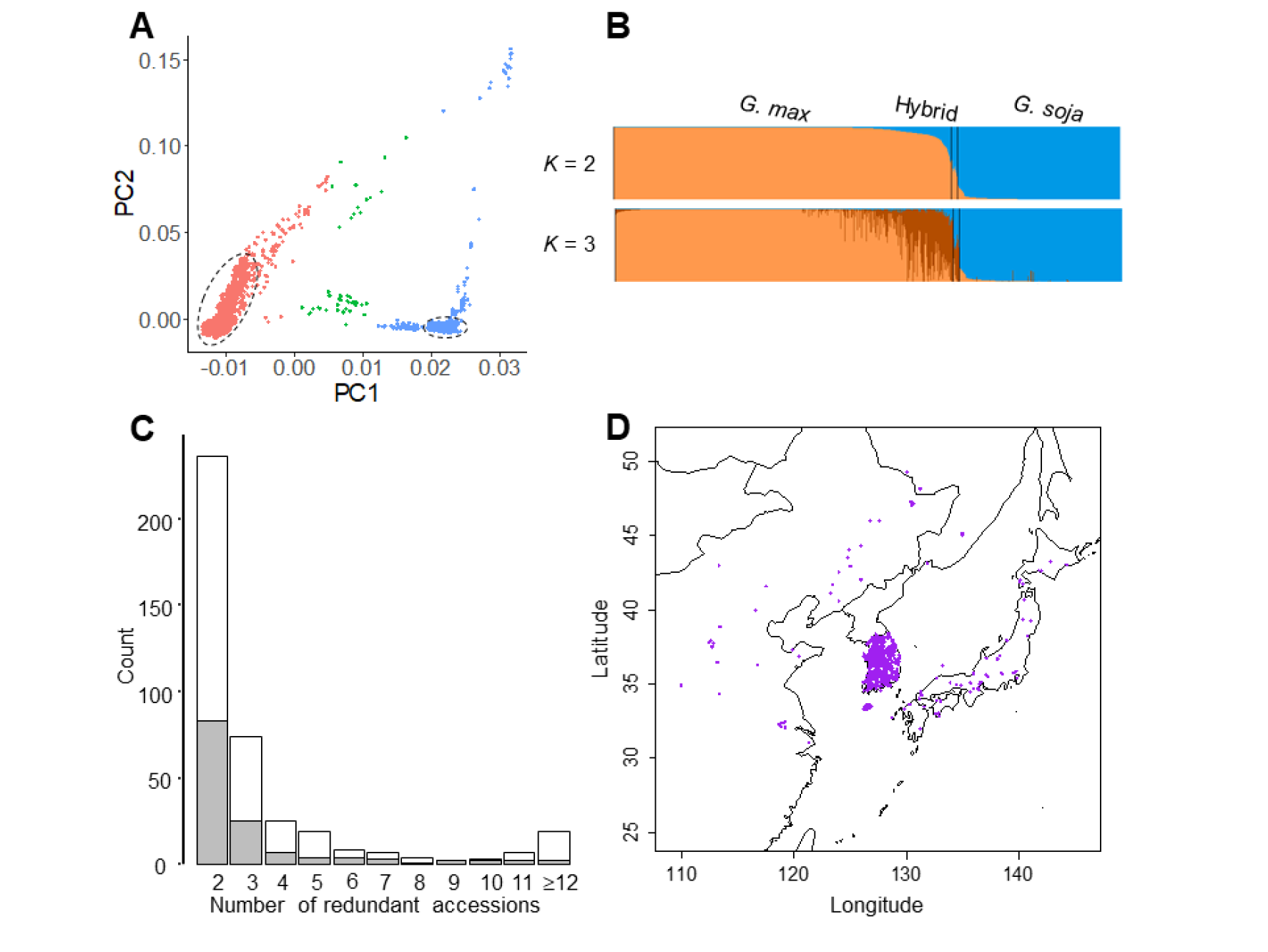
Population structure of the genotyped 4,234 soybean accessions. (A) Principal components of SNP variation. PC1 and PC2 indicate score of principal components 1 and 2, respectively. Each of PC1 and PC2 explained 15.6% and 2.7% of variance in the data. *Glycine max*, *G. soja*, and hybrids are shown by red, blue, and green dots, respectively. The majority of Korean accessions cluster together within dashed eclipses. (B) ADMIXTURE plots. The accessions were divided into three groups: *G. max*, *G. soja*, and their hybrids. (C) Distribution of number of redundant accession groups that showed <1.25% inconsistencies between the SNP calls. *G. max* and *G. soja* are shown by white and gray boxes. (D) Geographic distribution of the collection sites for *G. soja* accessions.

In our previous development and validation study (Lee et al. 2015), the SNP calls genotyped by the SoyaSNP array were highly reproducible, with inconsistencies of ≤1.17% observed within pairs of 27 duplicated samples after excluding missing genotypes in either sample. Several sets of near-isogenic isolines were genotyped (Table S3). Single-gene isolines (backcross-derived isolines for single genes) showed approximately 1.0% inconsistencies after excluding missing genotypes in either sample (e.g., 1.16% between Harosoy and L67-153 [Harosoy(6) x Higan]). As expected, a slightly higher level of inconsistency (up to 1.5%) was observed for samples from multiple-gene isolines (e.g. 1.48% between L62-667 [Harosoy(6) x T204] and OT94-51 [OT89-5/L71-802//OT89-6]). However, we occasionally observed that some soybean accessions that had no known pedigree relationship showed < 1.50% inconsistencies (e.g. 0.87% between Williams 82K and KLS85102). Therefore, we used a 1.25% inconsistency value as the cut-off to remove redundant or highly similar accessions from groups of duplicates or near isogenic lines. The same cut-off value was applied to filtration of wild soybean accessions. In each of the genotype duplicate sets, an accession with a sample call rate ≥99% was preferentially retained. For each of the near-isogenic line groups, the recurrent parent or representative single-gene isoline in case of the absence of parents was retained. Of the 4,184 accessions genotyped in this study, 1,148 (867 domesticated and 281 wild) were removed (Fig. 1C). The high rate of redundant and highly similar accessions has been frequently reported in worldwide germplasm collections (Food and Agriculture Organization of the United Nations 2010; McCouch et al. 2012). For domesticated soybeans, the major cause in the National Agrobiodiversity Center in Korea is probably the unknowing submission of the same accession with different collection sites and designators because there are many accessions with the same common name but with different collection sites or collectors. For wild soybeans, multiple accessions were collected from a narrow habitat area.

After the filtration, a final set of 3,036 genotyped accessions was available for population structure analysis (Table S1 and S2). Only a few accessions were removed from countries other than South Korea. As a result, overall proportion of soybean accessions among countries in this 3,036 set was similar to that of the 4,234 set (Table S2). In the 3,036 set, 1,957 were *G. max* accessions and 1,079 were *G. soja* accessions. Representative *G. soja* accessions from China, Russia, and Japan remained to evenly reflect the geographic distribution of native *G. soja* in each of these country regions (Fig. 1D).

### 3.2. Population structure

ADMIXTURE (Alexander et al. 2009) and PCA (Patterson et al. 2006) were used to infer population structure of the 3,036 non-redundant soybean set using 117,095 SNPs (heterozygous rate < 0.02 and minor allele frequency > 0.02). As observed in the analysis of our total population of 4,234 accessions, the 3,036 accessions were clearly divided into two large groups, representing *G. max* and *G. soja* (Fig. S1). Both the estimated cross-validation (CV) error plot from ADMIXTURE and scree plot from the PCA supported the presence of two large groups (Fig. S1), although the slopes did not level off, which is likely because of subgroupings within the two large groups. The clear separation of *G. max* and *G. soja* groups might be expected by the ascertainment bias, which favored selection of *G. max* SNPs (Lee et al. 2015), and the sampling bias. Unlike the previous observations that the genome diversity level was > 2-fold lower in the domesticated soybeans relative to that in the wild soybeans, the diversity level of the domesticated soybeans (mean per-site nucleotide diversity (π = 0.189) estimated from the 117,095 SNPs was ~1.58-fold lower than that of the wild soybeans (π = 0.298) and nearly two times more domesticated soybean accessions were used for the population structure analyses. The 3,036 set also contained excessive numbers of accessions collected in South Korea in both the domesticated and wild soybean groups. Interestingly, in the PCA space constructed with the first two PCs, *G. soja* accessions from Japan and China that were located at both ends of the *G. soja* cluster were almost equally close to the *G. max* cluster. However, in the PC1 and PC3 plot, accessions from Japan were closest to the *G. max* cluster and accessions from China were the most distantly related to the *G. max* cluster (Fig. S1).

When we analyzed *G. max* and *G. soja* separately, a somewhat distinct subpopulation structure was revealed. Within each of *G. max* and *G. soja* populations, CV errors of the ADMIXTURE runs decreased gradually without a steep drop (Fig. S2), whereas eigenvalues of the PCA runs showed steep decrease up to *K* = 5 (Fig. 2), indicating that there were at least four distinct subpopulations in each of the *G. max* and *G. soja* populations. Both the ADMIXTURE and PCA plots from the 1,957 domesticated soybeans did not show distinctive grouping (Fig. 2 and S2). The majority of South Korean domesticated accessions (~80%) formed a dense subpopulation likely because of recent overcollection. This notion is supported by that the rest of the Korea accessions were well mixed with Chinese, North Korean, and Japanese accessions, which did not show distinct subgrouping on a geographic basis. Notably, the North Korea accessions, which could be considered true landraces because of their collections during the first half of the 20^th^ century before modern breeding research, were evenly distributed across subpopulations. The improved cultivars were narrowly clustered in the PCA plot, indicating much lower diversity relative to that of the entire domesticated soybeans. The 1,079 wild soybean population showed distinctive subpopulations (Fig. 2 and S2), as shown in the analyses of the entire 4,234 population. The groupings were consistent with geographic distributions of the collection sites. Korean accessions and Japanese accessions formed a unique subpopulation, respectively. Chinese formed two subpopulations. Five accessions from the Russian border clustered together with those from northeast China. A strong relationship between subpopulations and their geographic distribution was notably exemplified by accessions from Jeju Island located 130 km off the southern coast of the Korean Peninsula; although they belong to the Korean subpopulation, all accessions from Jeju Island formed a subgroup that was the closest to the Japanese subpopulation (Fig. 2C).

**Figure 2.**
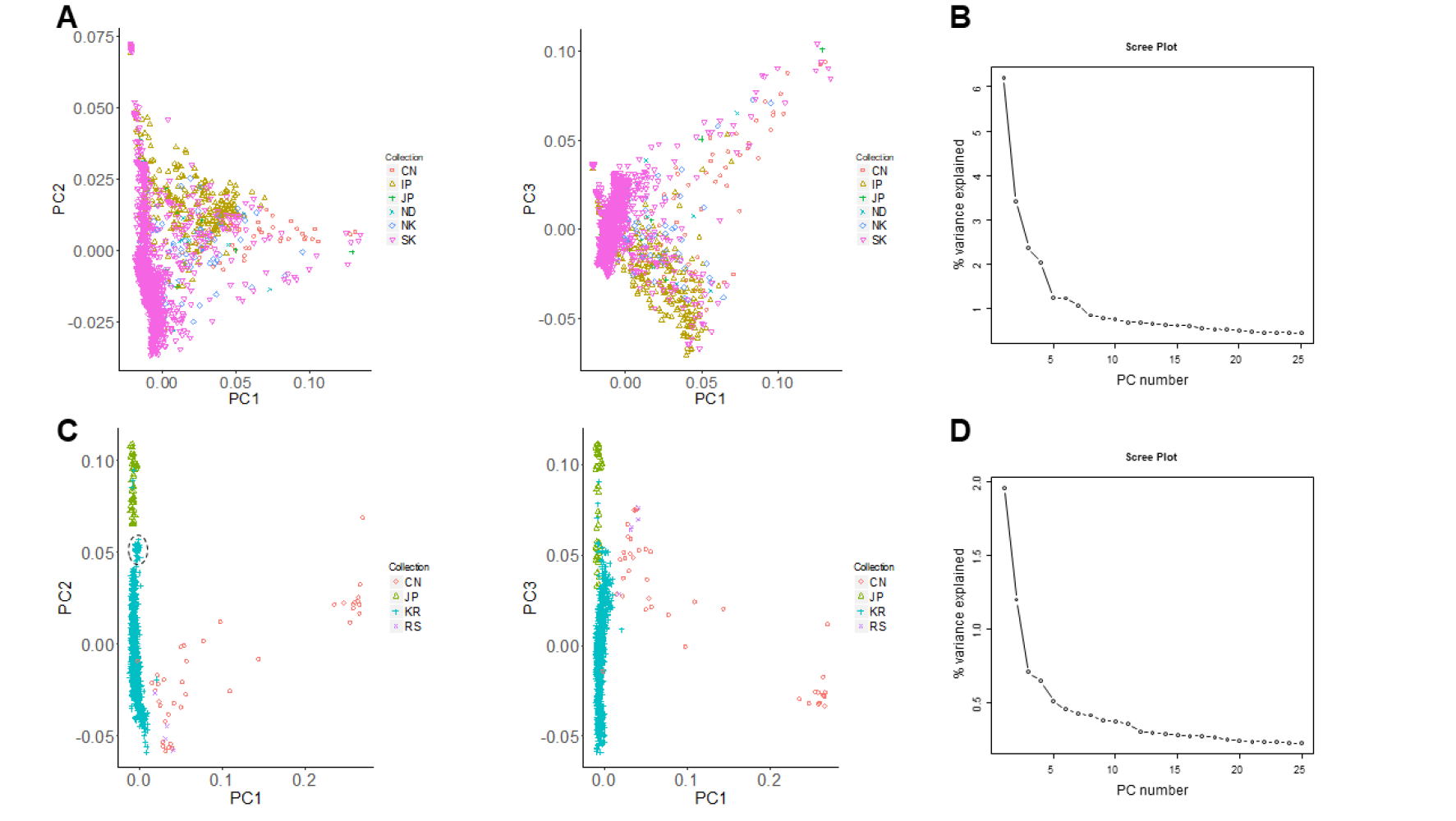
Population structures of 1,957 domesticated and 1,079 wild soybean accessions in the 3,036 non-redundant soybean accession set. (A) Principal components (PC) of SNP variation in the domesticated population. The plots show the first three principal components. The countries of collection or improvement status of the soybean accessions in (A) and (C) are represented by two-letter codes—CN, China; IP, improved breeding line; JP, Japan; ND, not determined; NK, North Korea; RS, Russia; and SK (KR), South Korea. (B) Scree plot of the PC number and their contribution to variance from principal component analysis of the domesticated accessions. (C) Principal components of SNP variation in the wild population. The plots show the first three principal components. A cluster of accessions from Jeju Island is indicated by a dashed eclipse. (D) Scree plot of the PC number and their contribution to variance from principal component analysis of the wild accessions.

### 3.3. Detection of SNPs associated with domestication history

Domestication is a process of continuous artificial selection of a group of traits, collectively called domestication syndrome. The domestication process has produced selective sweeps with significant reductions in nucleotide diversity (Doebley et al. 2006; Hufford et al. 2012; Chung et al. 2014) on limited regions of the genome (approximately 5 ~ 10% of the genome). Numerous recent whole-genome resequencing studies have effectively detected the selective sweeps, which are associated with domesticated genes (Meyer and Purugganan 2013), by examining reduction of diversity (ROD) in windows along chromosomes. However, our SoyaSNP array data are not dense enough to detect the reduction of diversity. Thus, we attempted to detect SNPs associated with domestication using a case-control GWA method that analyzed binary domestication phenotypes, which were determined by presence (*G. max*) or absence (*G. soja*) of domestication.

To test if our case-control GWA method enabled to find genes or chromosomal regions underlining binary phenotypes in our 3,036 non-redundant population, we chose two highly studied phenotypes—flower and seed-coat colors—which are monogenic and multigenic, respectively. Because our population was highly structured, we performed logistic regression model analysis conditional on a list of subpopulation-specific SNPs. The selected specific SNPs included one perfect domestication-specific SNP with each allele being perfectly correlated with *G. max* or *G. soja* membership of soybean accessions, and ten subpopulation-specific SNP within each of the *G. max* and *G. soja* populations (Table S4). The flower color phenotypes were divided into absence or presence of anthocyanin deposition colors. The seed-coat color phenotypes were divided into absence or presence of anthocyanin deposition colors. Using the conditional logistic regression model, we detected a broad and strong peak for flower color with the most significant SNP (max –log_10_*P* = 80.6) located at 17,877,234 on chromosome 13 (Figs. 3A, S3, and S4C). This peak area contained the *W1* locus, which is the major locus determining flower color (Zabala and Vodkin 2007). However, the most significant SNPs were located ~ 500 kb off the position of the *flavonoid 3*′*5*′*-hydroxylase* gene, which is the causal gene of the *W1* locus. To understand this region, we estimated pairwise LD for SNPs from 16.8 Mb to 18.8 Mb. A strong LD pattern was observed between all the SNPs under the most significant SNPs, however no clear LD pattern was observed near the *flavonoid 3*′*5*′*-hydroxylase* gene. Although the result might be due to a SNP density insufficient for a long LD block, it has been often observed that a causal gene for a strong peak are not always corresponding with the highest−log_10_ *P* value (Segura et al. 2012; Yano et al. 2016).

We detected more than 30 peaks exceeding a significant threshold (-log_10_ *P* ≥ 7.07) for seed-coat color (Figs. 3B, S3, and S4). The top three peaks were correlated with three known major loci (*I*, *R*, and *T* on chromosomes 8, 9, and 6, respectively) that control the deposition of various anthocyanin pigments in seed coat (Yang et al. 2010). The highest peak was generated from a chromosomal region surrounding the inverted CHS gene repeats, which is the causal region of the *I* locus (Clough et al. 2004). A strong LD pattern was observed at the *I* locus region. An SNP AX-90432942 on chromosome 6 with the second highest -log_10_ *P* value = 69.8 was generated from flavonoid 3’ hydroxylase, which is the causal gene of the *T* locus. Finally, the SNPs near the R2R3 MYB transcription factor gene, a strong candidate gene for the *R* locus, reported by Gillman et al. (2011) were not significantly associated with seed-coat colors. However, the gene is one of the R2R3 MYB transcription factor genes tandemly repeated in this chromosomal region and numerous highly significant SNPs were observed 100-kb away from the proposed *R* gene. Interestingly, the peak on chromosome 13 are located on the *W1* locus that influences seed-coat color in a case of the homozygous recessive *it* genotypes (Palmer et al. 2004). Considering such a wide range of soybean seed-coat color variations, the detection of numerous minor peaks is not surprising, as reported by Song et al. (2016), although it is still surprising to detect this large number of significant peaks using binary phenotyping data. Nevertheless, we think that some of those minor peaks are inevitably false because the limited number of conditional SNPs could not correct for all inflation of the statistic caused by population substructure.

**Figure 3.**
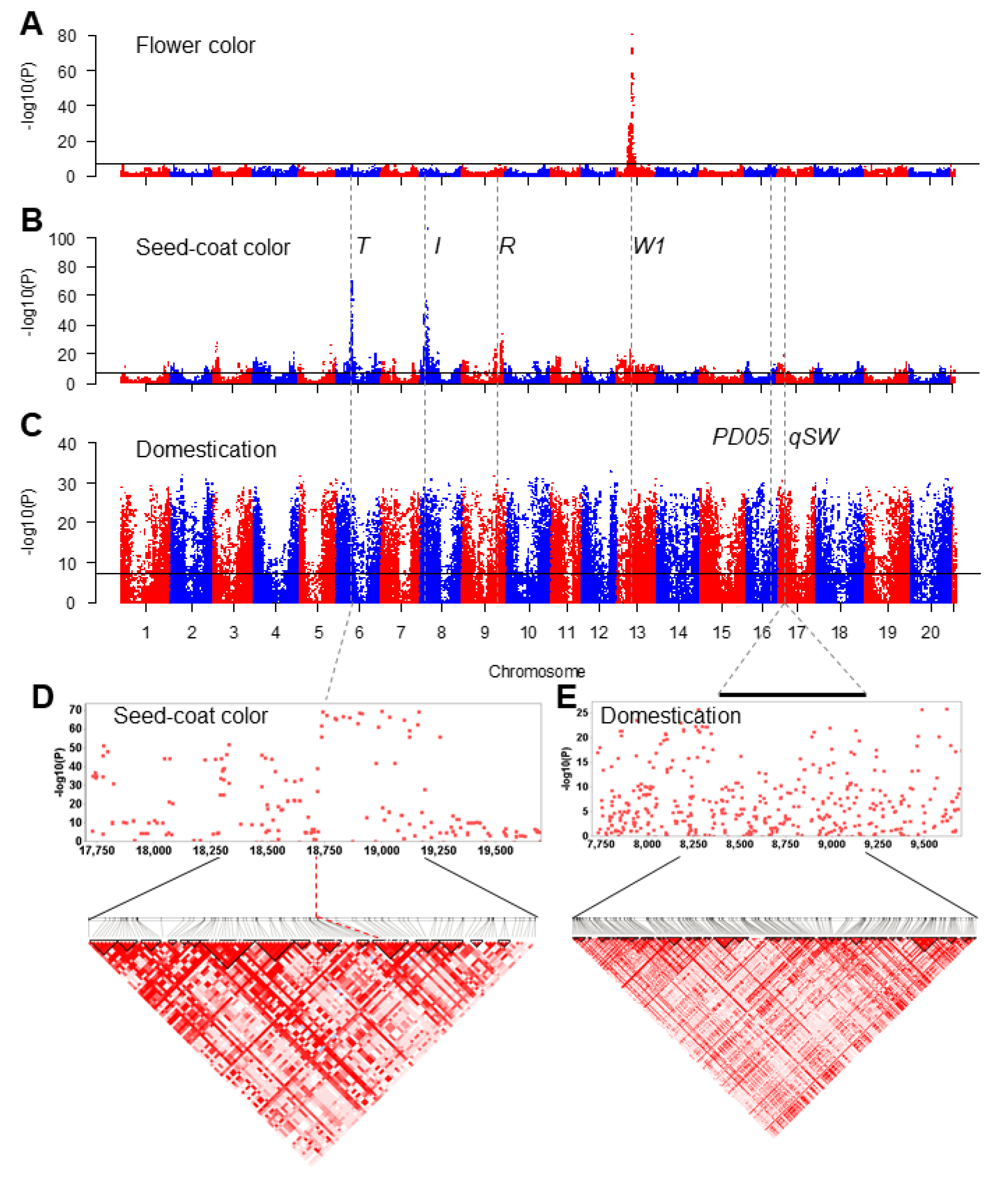
Genome-wide association scans for 3,036 soybean accessions for flower color, seed-coat color, and domestication. (A) Manhattan plot for flower color. The solid horizontal line denotes the Bonferroni-adjusted significance threshold. Chromosomal regions of known genes (*T*, *I*, *R*, *W1*) or loci (*PD05* and *qSW*) are indicated by dashed vertical lines. (B) Manhattan plot for seed-coat color. (C) Manhattan plot for domestication. (D) Local Manhattan plot (top) and LD heatmap (bottom) surrounding the *T* locus on chromosome 6. Dashed lines indicate the region of the *T* locus. Physical locations (kb) are indicated under the Manhattan plot. (E) Local Manhattan plot (top) and LD heatmap (bottom) surrounding the *qSW* locus on chromosome 17. A bar indicate the region of the *qSW* locus.

Since our logistic regression model could readily detect loci associated with flower and seed-coat colors using binary phenotypes, we performed GWAS for domestication syndrome using the binary domestication phenotypes. For this analysis, we excluded the perfect domestication-specific SNP in the list of subpopulation-specific SNPs (Table S4). We detected numerous peaks for domestication syndrome over the genome, as expected from the previous studies (Hufford et al. 2012; Meyer and Purugganan 2013; Chung et al. 2014) that showed that domestication features covered approximately ~7% of the crop genome. To examine if previously detected domestication regions were also detected in the current study, we compared peak locations from our study with selective signals previously detected for two domestication traits, pod dehiscence and seed weight (Figs. 3 and S4D), which are two of the most critical domestication traits among the traits assayed by Zhou et al. (2015). The 190-kb region (*PD05*) responsible for pod dehiscence was also detected in our study with three highly significant SNPs (-log_10_ *P* ≥ 20) and the *qSW* locus for seed weight was detected with > 30 highly significant SNPs (-log_10_ *P* ≥ 20). Lengths of strong LD blocks under the peaks corresponding to the *PD05* and *qSW* loci were not sufficient to define the selective sweeps that have been known to be > 100 kb, as observed in our LD analyses for the flower and seed-coat color loci. Flower and seed-coat colors analyzed in this study are considered domestication-or diversification-related morphological features because nearly all wild soybean accessions have purple flowers and black seed coats. As expected, their major loci, *W1*, *I*, *R*, and *T* were also detected with highly significant SNPs (-log_10_ *P* ≥ 20) in this GWAS for domestication syndrome (Fig. 3).

### 3.4. Extraction of a representative set

Our population structure and diversity analysis of the 3,036 non-redundant soybean population resulted in the identification of Japan as a likely center of soybean domestication. Since a fundamental assumption of model-based methods, such as ADMIXTRUE and PCA, is that the sample available for analysis is representative of the entire population distribution, sample sizes of subpopulations can substantially affect population stratification and ancestral population inference (McVean 2009; Shringarpure and Xing 2014). To investigate the possibility that excessive numbers of domesticated or Korean soybean accessions might have caused bias in inference of population structure of wild and domesticated soybeans, we obtained representative domesticated and wild soybean sets by selecting one from each of tightly distributed soybean miniclusters in the PCA plots, with a caution that overall distribution patterns are maintained (Fig. S5). For the representative set of wild soybeans, we filtered the population of the tightly distributed Korean accessions and selected 50 diverse Korean wild accessions (Table S2). In addition, four wild soybean anomalies misplaced to subpopulations different from subpopulations predicted by their collection site records were excluded; three Korean and one Chinese accessions (Fig. S6). For the representative set of domesticated soybeans, we selected 50 diverse *G. max* accessions that represent diversity of 1,957 *G. max* accessions. The results of the AMOVA indicated that the overall genetic structure observed in the 3,036 non-redundant soybean population was well represented by the extracted representative set with some decrease of genetic diversity in the *G. max* population (Table S5). PCA plots from the resultant representative set of 194 soybean accessions showed distribution patterns similar to those from the 3,036 non-redundant soybean accessions, although relative sizes of *G. max* and *G. soja* distributions in the PCA spaces constructed with the first and third PCs were reversed. The last drop of eigenvalues from the PCA runs occurred between *K* = 5 and *K* = 6 (Fig. S7), indicating that there were five distinct subpopulations.

### 3.5. Center of soybean domestication

Because *G. soja* can be found *in situ* across most of the East Asia, it is important to establish the population structure, if any, of a diverse collection of *G. soja* accessions and to associate one or more of these populations with a collection of domesticated *G. max* varieties. To perform these experiments, we analyzed the population structure of the representative set of 194 soybean accessions with ADMIXTURE, and found *K* = 5 populations based on the estimated CV error plot (Fig. S7). Thus, the soybean accessions were partitioned into one *G. max* (Gm) and four *G. soja* (Gs-I, Gs-II, Gs-III, and Gs-IV) subgroups (Fig. 4A, B). Wild accessions from China were divided into two subgroups, Gs-I and Gs-II. The Gs-I group showed the least diversity (Table 1) and most of them distributed in the middle region of the Yellow River basin. The Gs-II group was dispersed across northeast China, south China, and the Russian border of northeast China. Interestingly, this grouping result is remarkably similar to that obtained by a recent comprehensive study that showed that, by analyzing a total of 712 *G. soja* individuals from 40 natural populations in China, Chinese wild soybeans were grouped into two main subgroups, which were one from the Yellow River basin and the other from northeast China and south China (Guo et al. 2012). The Gs-IV distributed in Japan. The Gs-III showed the greatest diversity and distributed in South Korea and most of them appeared to be ancient admixture between Gs-II and Gs-IV. Interestingly, despite clear separation of the Chinese *G. soja*, diversity level of the combined population of Gs-I and Gs-II was similar to those of Korean or Japanese *G. soja*. An independent Gm group appeared from *K* = 2 to *K* = 5 (Fig. 4C). Interestingly, major genomic fractions of the Gm subgroup consistently appeared as minor genomic fractions of the Gs-IV and minor genomic fractions of the Gm group was a genomic fraction of Gs-I ancestry. The results suggested that after domestication of the Gm subgroup from the Gs-IV subgroup, the Gm subgroup was substantially diversified by introgression of the Gs-I genomic fractions. One of the Gs-I accessions, PI 459046, appeared to be a *G. max* x *G. soja* hybrid, although its genomic fraction (~22%) from *G. max* are lower than our hybrid filtration criteria (30% domesticated ancestry), which was less stringent than 20% domesticated ancestry used in other admixture studies (e.g.Wang et al. (2017)).

**Figure 4.**
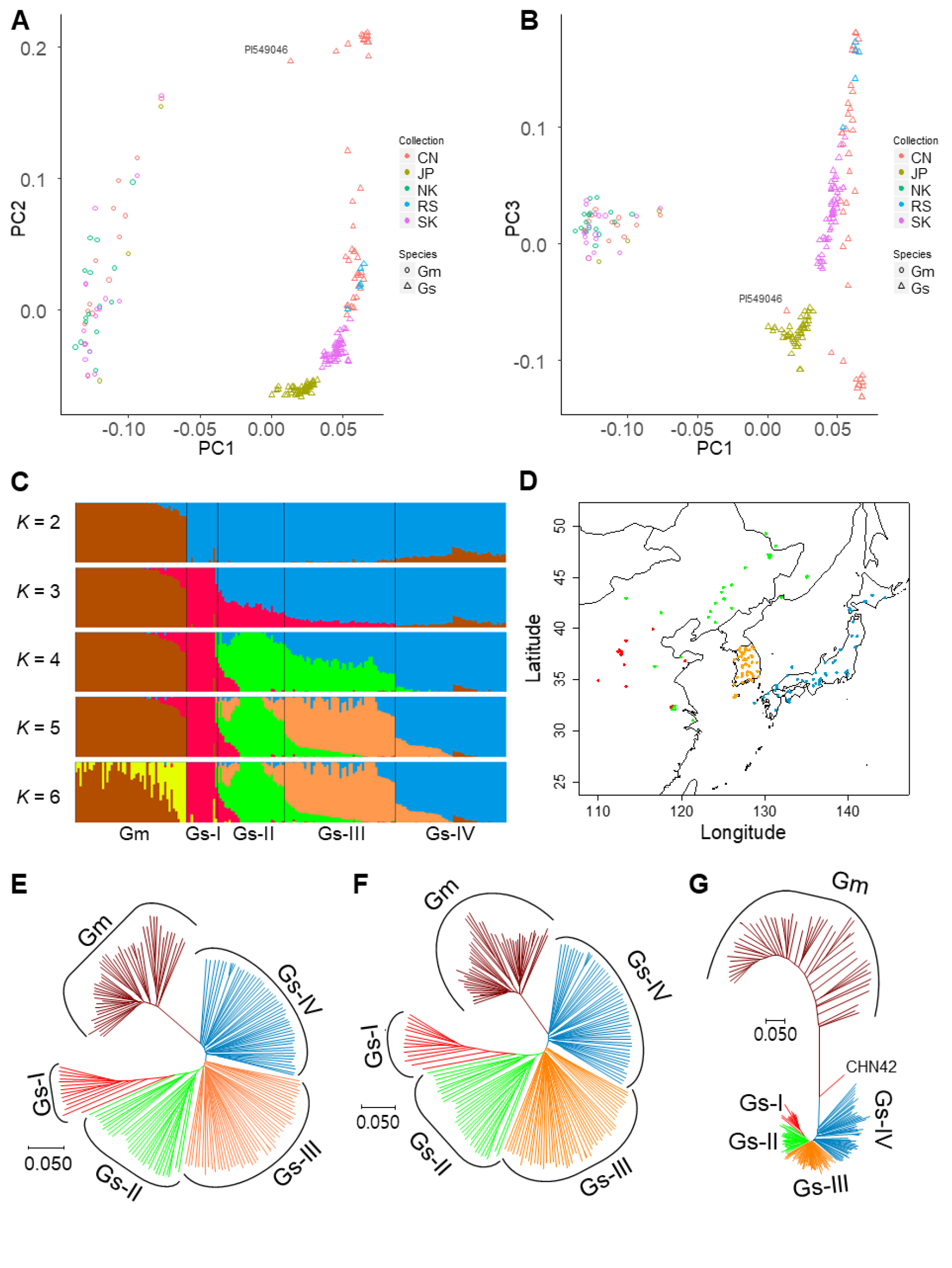
Identification of the domestication center of *G. max*. (A, B) Principal components plots of SNP variation. PC1, PC2, and PC3 indicate score of principal components 1, 2, and 3, respectively. Each of PC1, PC2, and PC3 explained 12.0%, 5.2%, and 2.6% of variance in the data. Countries of collection of the soybean accessions and species names are represented by two-letter codes—CN, China; JP, Japan; NK, North Korea; RS, Russia; SK, South Korea; Gm, *G. max*; and Gs, *G. soja*. A putative hybrid PI 459046 is labeled. (C) Population structure of 50 *G. max* (Gm) and 144 *G. soja* (Gs-I, Gs-II, Gs-III, and Gs-IV) accessions inferred using ADMIXTURE. Each color represents one population. PI 459046 showed ~20% of ancestral genomic fractions from *G. max*. (D) Geographic distribution of the four *G. soja* subgroups. Gs-I is red, Gs-II green, Gs-III orange, and Gs-IV blue. (E, F, G) Neighbor-joining phylogenetic tree of 194 soybean accessions based on the SNPs genotyped by the 180K AXIOM SoyaSNP array, with evolutionary distances measured by the *p* distance. The taxa used in the neighbor-joining tree and bootstrap values from 1000 bootstrap replications at branches are described in Fig. S8. (E) Phylogenetic tree based on 117,095 SNPs. (F) Phylogenetic tree based on 108,897 SNPs, which are weakly or not significantly associated with domestication traits. (G) Phylogenetic tree based on 8197, which are very significantly associated with domestication traits. PI 459046 from group Gs-I clusters between Gm and Gs-IV likely because of contribution of ancestral genomic fraction from Gm.

We constructed a neighbor-joining (NJ) tree for the representative soybean set (Fig. 4E and S8). The tree showed that all *G. max* accessions formed a monophyletic cluster. Although *G. max* was artificially selected recently, terminal branch lengths were similar to those of *G. soja* likely because of ascertainment bias that more SNPs were selected from *G. max* than from *G. soja* (Lee et al. 2015). The population of the nearest branches, which were basal to the *G. max* soybean lineage, was *G. soja* subgroup Gs-IV. Within the Gs-IV that contains wild soybeans from Japan, those from eastern Japan area were closer to the *G. max* soybean lineage. To measure population differences and similarities, we calculated the fixation index values (*F_ST_*) (Holsinger and Weir 2009) between *G. max* and each *G. soja* population (Table 2). The pairwise *F_ST_* values ranged from 0.201 to 0.334. The value of *F_ST_* for the *G. max* and Gs-IV populations was the smallest, suggesting that *G. max* was domesticated directly from *G. soja* subpopulation Gs-IV. The level of population differentiation among *G. soja* subpopulation was much lower than that between *G. soja* and *G. max*, similar to the case of rice (Huang et al. 2012). However, *F_ST_* values between *G. soja* subpopulations corresponded with their geographic distances. The regions containing those wild populations that are phylogenetically close with cultivars could be proposed as the domestication region of crops (Matsuoka et al. 2002; Spooner et al. 2005). Thus, our results suggested that soybeans had been most likely domesticated only once in eastern Japan.

**Table 1.** Mean per-site nucleotide diversity (*π*) of *Glycine max* (Gm) and each of *G. soja* groups (Gs-I to Gs-IV) in the representative soybean set

| Group | Representative soybean set |  | Representative soybean set and 45 wild soybeans from Zhou et al. (2015) |  |
| --- | --- | --- | --- | --- |
| | Number of accessions | $\pi$ | Number of accessions | $\pi$ |
| Gm | 50 | 0.236 | 50 | 0.237 |
| Gs-I | 14 | 0.198 | 29 | 0.222 |
| Gs-II | 30 | 0.276 | 45 | 0.283 |
| Gs-I and Gs-II | 44 | 0.290 | 74 | 0.301 |
| Gs-III | 50 | 0.294 | 57 | 0.299 |
| Gs-IV | 50 | 0.276 | 58 | 0.283 |
| Gs-III and Gs-IV | 100 | 0.296 | 115 | 0.302 |

Because the number of *G. soja* accessions from China in our representative set was smaller than those from Korea and Japan, the population structure revealed by our representative set was further resolved bv incorporating the SNP data from 62 *G. soja* accession genomes resequenced by Zhou et al. (2015). By intersecting these SNPs with the set of 117,095 high-quality SNPs selected in this study, we extracted 103,801 SNPs, which were shared between the genome resequencing data and our representative set. Of the 62 accessions, only 45 were incorporated into the representative set because of the high level (eleven accessions, > 20%) of heterozygous SNPs, hybrid (one), and overlapping (five) (Fig. S9 and Table S6). The resultant expanded set contained twelve diverse accessions from Zhejiang, China and one accession from Taiwan, thus increasing geographic coverage of this study further down to southern China. In results, the diversity level of *G. soja* accessions from China and its Russian border was similar to those from Korea or Japan (Table 1). Population structure of the expanded set inferred from ADMIXTURE and PCA was quite similar to that of our representative set, although the *G. max* accessions appeared to be divided into two groups likely because of the high level of heterozygous SNPs from the genome resequencing data (Fig. S10, S11, and Table 2). Phylogenetic analysis and estimated *F_ST_* values between subpopulations indicated that *G. soja* accessions collected from eastern Japan were closest to the *G. max* soybean lineage.

**Table 2.** *F_ST_* values between *Glycine max* (Gm) and each of *G. soja* groups (Gs-I to Gs-IV) and between *G. soja* groups

| Representative soybean set |  |  |  |  |
| --- | --- | --- | --- | --- |
|  | Gs-I | Gs-II | Gs-III | Gs-IV |
| Gm | 0.334 | 0.267 | 0.226 | 0.201 |
| Gs-I |  | 0.160 | 0.180 | 0.214 |
| Gs-II |  |  | 0.062 | 0.113 |
| Gs-III |  |  |  | 0.058 |
| Representative soybean set and 45 wild soybeans from Zhou et al. (2015) |  |  |  |  |
| Gm | 0.323 | 0.264 | 0.227 | 0.204 |
| Gs-I |  | 0.164 | 0.176 | 0.209 |
| Gs-II |  |  | 0.066 | 0.118 |
| Gs-III |  |  |  | 0.055 |

To evaluate the distribution of SNPs associated with domestication syndrome across soybean subpopulations, we divided the 117,095 SNPs into 8,197 SNPs highly significantly associated with domestication traits and 108,898 SNPs weakly or not significantly associated with domestication traits. The 8,197 SNPs (-log_10_ *P* > 17) were selected because the previous studies have shown that ~7% of the crop genome is domestication-related (Hufford et al. 2012; Xu et al. 2012; Chung et al. 2014). Our GWAS also indicated that, in our GWAS population, strong LD extent under the highly significant SNPs in a peak tended to be much shorter than chromosomal extent under all the significant SNPs of the peak. The tree constructed from the 108,898 SNPs (Fig. 4F) was similar to that constructed from the 117,095 SNPs in their overall grouping and branch patterns, except that the branch length and grouping of the *G. max* clade were slightly different to each other. Grouping patterns in the tree constructed from the 8,196 SNPs (Fig. 4G) were similar to those in the trees constructed from both 108,898 SNPs and 117,095 SNPs, except that the putative hybrid PI 459046 moved the closest to the *G. max* clade. Interestingly, the lengths of the basal and terminal branches for the *G. max* clade and the *G. soja* Gs-IV clade in the tree from the 8,196 SNPs became distinctively longer than those in the other *G. soja* clades. The results indicated that initial major artificial selection for soybean domestication was limited to a *G. soja* group from Japan.

## 4. Discussion

The present study analyzed genome-wide SNP variations obtained from thousands of soybean accessions, the majority of which were collected from the Korean peninsula. The results provide insight into the development of strategies for efficient and directed germplasm use as well as for collection of novel landraces and wild relatives. Population structure and grouping analyses revealed strong correlations between genetic distance and geographic distance in *G. soja* (wild soybean) populations and weak correlations in *G. max* (domesticated soybean) populations. *G. soja* accessions were divided into four distinct subgroups; Gs-I and Gs-II from China and its Russian border, Gs-III from Korea, and Gs-IV from Japan. The results suggest that although the Korean territory is much smaller than Chinese and Japanese territories, the ocean-imposed geographic separation among these countries has been a major contributor to the evolutionary divergence of *G. soja*. Most of the Gs-III group from Korea appeared to be ancient admixtures between Gs-II and Gs-IV, suggesting that *G. soja* spread from each of China and Japan might be mixed in Korea. Thus, Korean wild soybeans are more valuable resources than the other countries’ wild soybeans because they alone provide variations from two large subgroups. Interestingly, accessions from Jeju Island off the southern coast of the Korean Peninsula are the closest grouped to the Gs-IV group among members of the Gs-III group, indicating that although our estimated *F_ST_* values between *G. soja* groups denied appearance of a new distinct group, more extensive sampling from diverse areas will likely reveal better correlations between geographic and genetic distances among *G. soja* subpopulations. The *G. max* population was divided into four subgroups. However, it was apparent that the subgrouping did not reflect geographic origins. Particularly, landraces from North Korea that would be considered true landraces based on their collection time appeared in every subgroup. The majority of South Korean landraces that had been collected recently were grouped together. The majority of improved accessions from the United States were clustered closely together, supporting a previous observation (Hyten et al. 2006). Taken together, our results suggest that while *G. soja* germplasm will require additional sampling from diverse indigenous areas to expand the germplasm base, *G. max* germplasm is saturated in terms of genetic diversity. Thus, extensive genotyping and phenotyping of extant *G. max* germplasm would be the next step to expand the germplasm base of *G. max*.

Our results provided strong support for a single origin of *G. max* from eastern region in Japan, although pointing to a specific region in Japan likely requires analysis of more extensive wild and landrace soybean accessions from Japan. Whether a crop species stems from a single domestication event or from multiple independent domestications has been consistent with whether the domesticated species are monophyletic or polyphyletic, respectively, in the phylogenetic trees constructed from both the domesticated and wild progenitor species. Although diversity of chloroplast DNA, which represents maternal lineage of soybean, revealed multiple lineages of domesticated soybeans, analyses of recent genome-wide soybean variation data (Guo et al. 2010; Lam et al. 2010; Chung et al. 2014; Zhou et al. 2015) consistently showed the monophyletic nature of *G. max*, as observed in this study. In other words, recent soybean phylogenetic studies collectively indicated a single origin of *G. max*. The best examples of monophyletic grouping are wheat and barley, which appear to have been domesticated once from their wild ancestors in the Fertile Crescent (Badr et al. 2000; Ozkan et al. 2002). The origin of barley was further supported by the genome sequences of five 6,000-year-old barley grains (Mascher et al. 2016). In cases of rice and common bean that showed polyphyletic groupings, single or multiple regions of origin of these crop species are still contentious (Molina et al. 2011; Bitocchi et al. 2012; Huang et al. 2012). This controversy may have arisen because most modern wild accessions studied represent descendants of ancient feralization of admixed accessions that resulted from hybridization events between domesticated species and wild species populations after domestication (Wang et al. 2017), indicating that one of the previously thought independent origin regions might be a secondary origin region. The grouping of PI 459046 in this study is a good example that shows how hybrids could mislead inference of relationship between wild and domesticated crop species.

A recent comprehensive study of the archaeological records for soybean from Japan, China, and Korea indicated that Japan could have been a source of a large-seeded landrace of domesticated soybean that spread to Korea and subsequently to China (Lee et al. 2011). The archaeological records suggest that selection of large seed sizes occurred in Japan (Lee et al. 2011; Nakayama 2015) by 5,500 calibrated years (cal) before present (BP) and in Korea (Lee et al. 2011) by 3,500 cal BP. Seed size is clearly a domestication trait because the seed size of *G. soja* is much smaller than that of *G. max* landraces (Broich and Palmer 1980). However, the archaeological data were interpreted to suggest the multiple origins hypothesis of soybean. One particular reason is that the excavated tiny seeds were as old as 9,000–8,600 cal BP in northern China and 7,000 cal BP in Japan. However, the size of the seeds is similar to that of the seeds of present-day wild soybeans, and so would have been quite different from the landraces already grown in China by 2,500 BP. Another reason is that the interpretation was greatly influenced by a previous report that diversity of chloroplast DNA SSRs in wild and domesticated soybeans showed evidence for multiple origins of domesticated soybean (Xu et al. 2002). However, as mentioned above, numerous recent genome-wide soybean genome variation studies consistently show a single origin of *G. max*.

One of the main reasons that the previous studies pointed different regions in China as the center of soybean domestication is likely sampling bias. Our results suggested that wild accessions from China had genetic diversity level almost equal to those from Korea or Japan. However, most previous studies tended to neglect this fact. In an extreme case (Han et al. 2016), no accession from Korea and Japan was used, with the conclusion that central China is the initial domestication region. Another confounding factor is the inclusion of hybrid soybeans from natural mating between *G. soja* and *G. max*. Hybrid soybeans were not recognized in many previous soybean population studies, although hybrids between wild and domesticated species have been increasingly regarded as a major problem in studies of crop domestication history (Bitocchi et al. 2012; Wang et al. 2017). Furthermore, it was often assumed that a region in China is a center of soybean domestication because hybrid soybeans are frequently found in China (Han et al. 2016). However, of the 50 hybrids that we removed to avoid their potential confounding effects in this analysis, the majority (36 of 50) were accessions from Korea. The domestication of domesticated plant species from their wild ancestors arose from rapid evolutionary changes in the past 13,000 years of Holocene human history (Diamond 2002; Larson et al. 2014). The list of origins and the list of the most productive areas of most of major crops in the modern world are almost mutually exclusive. This could be explained by that the domestication origin of a crop was merely a region to which the most numerous and most valuable domesticable wild plant species were native. In this respect, our result that shows Japan as the domestication origin of soybean is not totally unexpected one.

Expanding on previous studies that reported genome-wide polymorphism data of soybean germplasm (Lam et al. 2010; Chung et al. 2014; Bandillo et al. 2015; Zhou et al. 2015; Valliyodan et al. 2016; Wang et al. 2016), our results show that samples intensively collected from Korea, which is a small area of the entire soybean distribution, provide sufficient amounts of data to underpin genome-wide population genetic questions that have been neglected or misled in the context of diversity and domestication panels of extant individuals. Our analysis demonstrates the value of current germplasm collections and how to expand the germplasm base. Furthermore, the findings show that a single major domestication event had occurred in a region of Japan. In addition, the high-density SNP array data enabled detection of domestication-associated SNPs and regions controlling important agronomic traits in a highly accurate manner. This suggests that our results will likely be useful for marker-assisted selection and genomic prediction to utilize unexplored genetic diversity in the soybean germplasm.

## Data accessibility

SNP genotype data are listed in Table S1 and are publicly available at Korean Soya Base (http://koreansoyabase.org/Data_Resource/).

## Acknowledgments

We thank Dr. Changyong Lee at Kongju National University for helpful comments in statistical analysis. We are grateful to Prof. Michael Gore at Cornell University for his critical reading of the revised version of this paper.

## Supplementary data

Supplementary data are available at DNARES online.

